# Resource competition affects the developmental outcome of the acoustic parasitoid fly *Ormia ochracea*

**DOI:** 10.1101/2024.04.25.590943

**Authors:** Jimena A. Dominguez, Brendan Latham, Laura C. Mongui, Addie Rossinow, Yeng Xiong, Briella V. Schmidt, Quang Vu, Blanca L. Torres-Lopez, Parker A. Henderson, Andrew C. Mason, Norman Lee

## Abstract

*Ormia ochracea* is an acoustic parasitoid fly where the adults are free-living, but their larval young depend on nutritional resources within host crickets for growth and development. In nature, gravid female flies rely on their ability to recognize and localize cricket calling songs to find suitable host species to parasitize. In depth investigations of fly behavior and the mechanistic bases of auditory perception require a reliable approach to propagate stable fly colonies in the laboratory. Previous work has demonstrated that flies can be propagated using a number of natural host cricket species, as well as cricket species that flies do not parasitize in nature. However, we lack a complete understanding of fly developmental outcomes when non-host cricket species are utilized to propagate fly colonies. In this study, we document the feasibility of using commercially supplied *Acheta domesticus* as a host species. We specifically test the hypothesis that host size and resource competition can affect developmental outcomes of *O. ochracea*. We performed manual parasitizations on crickets that varied in size, and resource competition was varied by manipulating the number of larvae used to parasitize a host cricket. A series of morphometric analyses were conducted on host crickets, and developmental outcomes were measured in terms of pupation success and eclosion success, pupal width, and eclosed adult fly size. In the absence of resource competition, we found that host cricket size did not affect pupation or eclosion success. In the presence of resource competition between two developing larvae within a host cricket, pupation and eclosion successes were impacted negatively, and the developing pupae were more likely to be smaller. These results confirm that resource competition among developing parasitoids can negatively affect developmental outcomes, and *Acheta domesticus* can be used effectively to propagate colonies of *O. ochracea* in the laboratory.

**Highlights:**

- The acoustic parasitoid fly *Ormia ochracea* can successfully develop within the house cricket *Acheta domesticus*.
- Pupal size and eclosed adult fly size varies positively with the size of host cricket.
- Resource competition negatively affects pupation and eclosion success.
- Resource competition resulted in smaller fly pupae that were less likely to eclose.

**Graphical Abstract:** 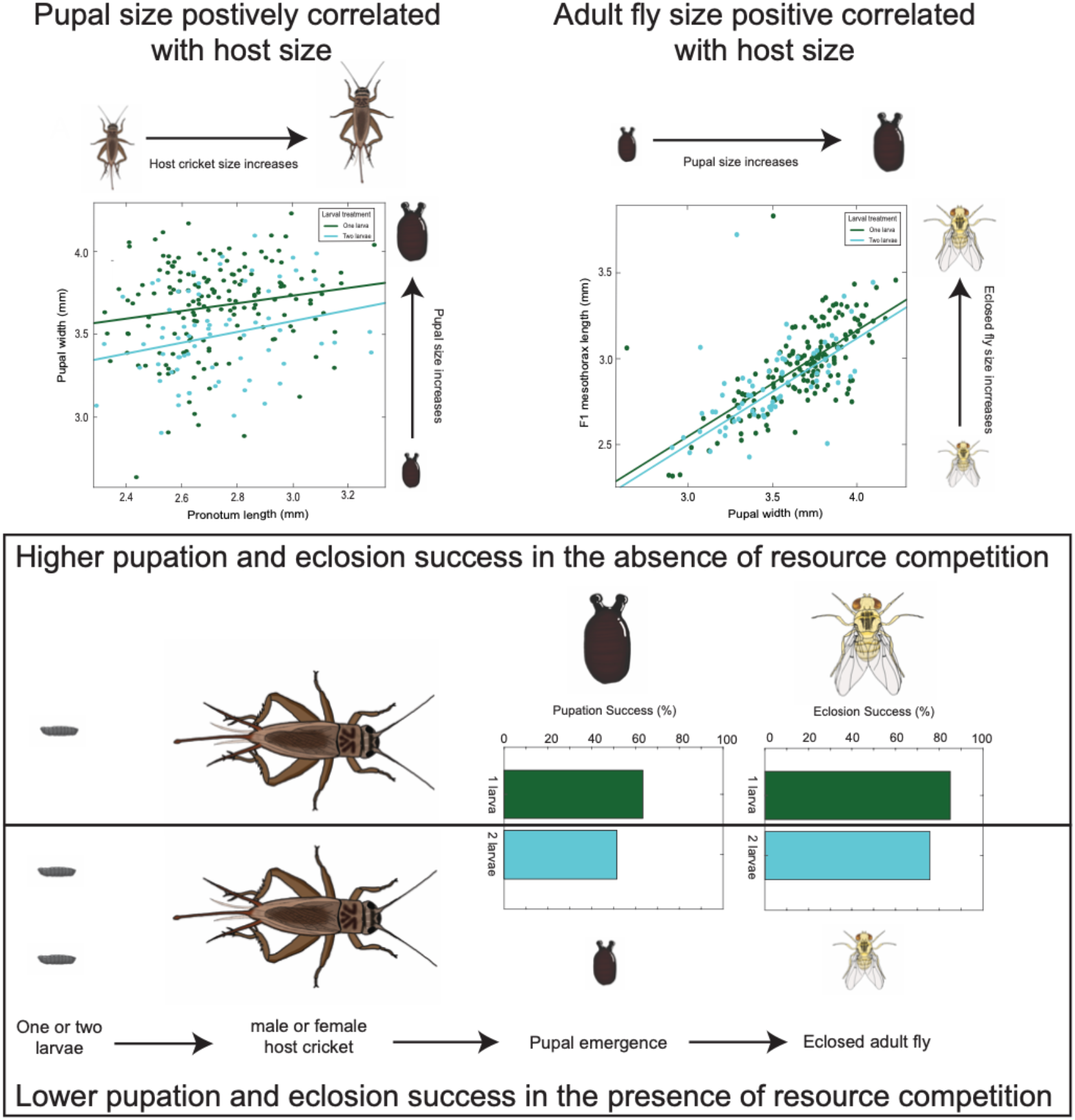

## Introduction

In insect parasitoids, competition for limited resources can affect trait expression and life-history evolution (Ode et al., 2022). While adult parasitoids are free-living, their larval young depend on nutritional resources within hosts for growth and development (Godfray, 1994). Consequently, the dynamics of resource availability, together with extrinsic (competition among adults for hosts) and intrinsic (competition among immatures) competition, can profoundly shape developmental trajectories and fitness outcomes (Harvey et al., 2013; Ode et al., 2022). For example, some parasitoids exhibit preferences for larger hosts and may parasitize such hosts with more eggs, as larger hosts may support the development of a larger number of offspring (Nechols and Kikuchi, 1985). Furthermore, emerging parasitoids’ body size is often positively correlated with host size (Cohen et al., 2005; Kouamé and Mackauer, 1991). Thus, the size of a host and level of competition for resources may affect resource availability and developmental outcomes of parasitoids.

The tachinid fly *Ormia ochracea* (Bigot 1889) is a larvi- and viviparous endoparasitic dipteran acoustic parasitoid that relies on host crickets for the development of their larval young (Cade, 1975; Godfray, 1994). Gravid females locate host crickets by homing in on cricket calling songs (Cade, 1975; Mason et al., 2001). Their auditory system is tuned to the dominant frequency of cricket calling songs (Oshinsky and Hoy, 2002; Robert et al., 1998, 1992), and song recognition is based on processing the temporal patterning of sound pulses (Jirik et al., 2023; Lee et al., 2019; Lee and Mason, 2017; Walker, 1993). Once *O. ochracea* detect suitable host songs (Gray et al., 2019, 2007), they engage in flying phonotaxis, land, and continue with walking phonotaxis to the sound source (Lee et al., 2009), where they deposit planidia (1st instar larvae) near or on top of host crickets. These planidia will wave their anterior end in the air and attach to a potential host cricket, and subsequently burrow into the host for development (Adamo et al., 1995a).

Larval development within a host cricket occurs for 7 to 10 days (Adamo et al., 1995a). During the first three days of development, the planidia appear to feed on hemolymph while leaving muscle tissue undisturbed. By the fourth day, planidia migrate to the abdomen, molt, attach to the abdominal wall, and form a respiratory funnel, which provides protection from the host immune system and allows the planidia to maintain contact with outside air. The planidia will continue to molt and feed on surrounding tissue. During the last two days of parasitization, planidia will feed on the fat body and abdominal and thoracic muscles, but will spare the digestive system and the central nervous system (Adamo et al., 1995a). Right before emergence from the host, larvae will purge their gut contents inside the host, which ultimately leads to cricket death. The larvae will then exit the host cricket, and pupate within a few hours.

In the field, flies can deposit one or several larvae on a host cricket (Adamo et al., 1995b). Both male and female crickets have been found to be parasitized by *O. ochracea* (Walker and Wineriter, 1991), but since *O. ochracea* are attracted to calling songs that are only produced by males, males have been found to be parasitized at a higher rate than compared to female crickets (Adamo et al., 1995b; Zuk et al., 1993). In the field, host crickets found to be parasitized by *O. ochracea* were most often found to be harboring 1 to 2 (Adamo et al., 1995b; Kolluru and Zuk, 2001) or more larvae (Gallagher et al., 2024), but it has not been reported in these studies whether some or all of these larvae are able to successfully develop to eclosion. When larviposition behavior was observed in the lab, crickets were never found to be parasitized with more than two larvae. However, cricket grooming behavior can help remove larvae before they burrow into the cricket (Vincent and Bertram, 2010a).

While previous work has mainly examined parasitization outcomes in *O. ochracea*’s natural host species (Adamo et al., 1995a; Broder et al., 2023; Cade, 1975; Thomson et al., 2012), few studies have looked at developmental outcomes in the house cricket *Acheta domesticus*, a cricket species that can be easily obtained from commercial suppliers. The ease of access to large supplies of *A. domesticus* can allow for the propagation of thriving laboratory colonies of *O. ochracea*. In this study, we investigate the effectiveness of using the house cricket *Acheta domesticus* as a host species to rear laboratory colonies of *Ormia ochracea*. We perform manual parasitizations on male and female crickets that vary in size, and experimentally manipulate the larval load per host to vary the level of resource competition. Our goals are to determine how host size and level of competition affect larval developmental outcomes, including pupation success, the resulting pupal size, eclosion success, and F1 adult fly size. If resource competition and size of the host affect larval developmental outcomes, we expect that higher levels of competition within smaller host crickets (typically males), will result in less ideal larval developmental outcomes.

## Methods

### Animals

Recently molted adult *Acheta domesticus* crickets (4-6 days post final molt) were acquired from a supplier (Bug Co., Ham Lake, MN). A total of 600 crickets (300 males and 300 females) were used as host crickets for the study. Ten gravid female *Ormia ochracea* from a laboratory colony that originated from Gainesville, FL were used as larval donors.

### Animal Care

#### Crickets

*A. domesticus* were kept at approximately 21°C over the course of the experiment. They were housed individually in 3.25 oz sauce cups with perforated lids to allow for respiration. We provided the animals with water and food (Purina Complete alfalfa rabbit feed pellets) *ad libitum*. Containers were checked daily for fly larval emergence, pupation, and animal deaths.

#### Flies

Donor *O. ochracea* were reared in temperature-, humidity-, and light-controlled environmental chambers (Power Scientific Inc, model DROS52503, Pipersville, PA) set to a 12 hr light / 12 hr dark cycle at 75% humidity, and provided with butterfly nectar (The Birding Company, MA) *ad libitum*. Eclosed F1 flies were kept in the same sauce cup container as the cricket they parasitized. Flies were fed butterfly nectar via a cotton stick that was changed daily.

#### Morphometrics

Crickets were prepared for imaging by cold anesthetization in a -20°C freezer for 10 minutes. This allowed the experimenter to orient and align the cricket dorsal aspect upward for imaging the pronotum (Fig. 1A) and the ventral aspect upward for imaging the femurs of the hind legs (Fig. 1B). These images were taken prior to the manual parasitization procedure. After taking the images, crickets were weighed using a digital scale (Sartorius Entris, 1241-IS Balance 120g x 0.1mg), given a unique cricket ID, and entered in our data collection sheet.

**Figure 1.**
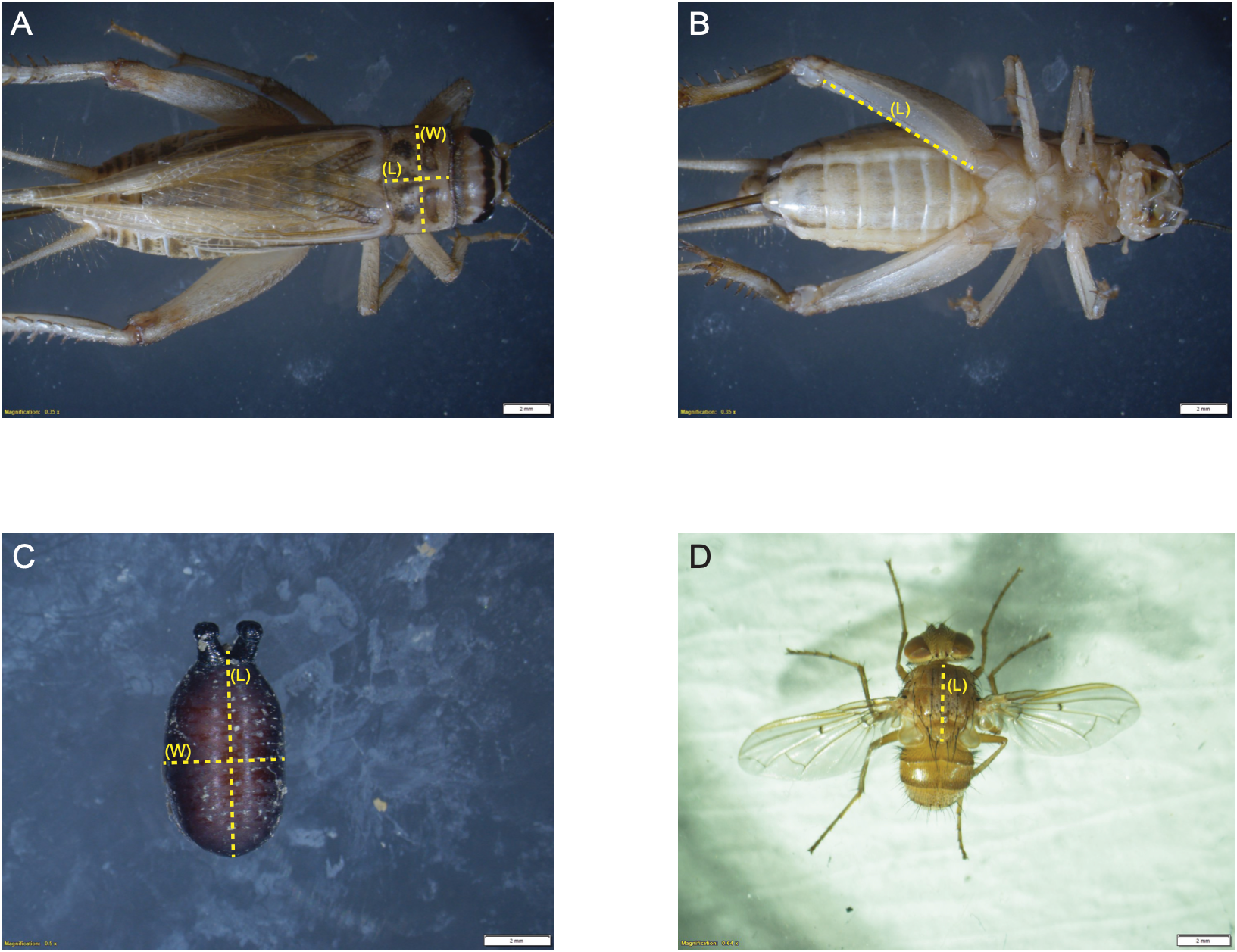
Cricket and pupal morphometrics. **(A)** Dorsal aspect of a female *Acheta domesticus*. Measurements of the length (L) and width of the pronotum (W) indicated by the yellow dotted lines. **(B)** Ventral aspect of the same female cricket, with a yellow dotted line indicating femur length (L) measurements. **(C)** A pupa with yellow dotted lines indicating length (L) and width (W) measurements. All images include a calibrated scale bar used to calibrate x and y pixel values in ImageJ for measurements. **(D)** Dorsal aspect of an eclosed F1 female *O. ochracea*. Measurement of the length (L) of the mesothorax indicated by a yellow dotted line. Scale bars in all panels represent 2 mm length.

Pupal images were taken before flies eclosed. This was done by orienting the pupae with the spiracles facing upward to allow for measuring of pupal length and width (Fig. 1C).

Images of crickets, pupae, and flies, were captured using Cell Sens Dimension (ver 3.24) interfaced with an Olympus SZX16 stereomicroscope equipped with a SDFPLAP0 0.8x objective and a DP80 Digital Camera. The images produced by this setup included a calibrated scale that facilitated measurements of morphology features (Fig. 1A-C). Captured images were imported into ImageJ (ver 1.53) for morphometric measurements. The scale bar in the image was used to calibrate pixel values to real-world size measurements. We then used the line tool to measure the pronotum length and width, and the femur length in crickets. In flies, we used this tool to measure pupal length and width (Fig. 1C), and mesothorax length (Fig. 1D).

#### Parasitizations

Manual parasitizations were performed by extracting larvae from 10 freshly dispatched donor flies. This involved dissecting the abdomen of a fly and spreading out larvae contained in the fly reproductive tract on a petri dish lined with filter paper. Freshly extracted larvae move around in the petri dish and often stand on their posterior ends and move its anterior end in a wave-like motion. On a piece of damp filter paper, larvae are most active within the first two hours after removal from the gravid fly, but can potentially survive for 7-8 hours (Beckers et al., 2011). In this state, larvae can easily attach to a wooden probe. Once attached to the probe, we placed individual larva on top of the articular sclerite located just above the anterior margin of the cricket’s thorax and directly underneath the pronotum. Larvae from each of the donor flies were used to infest 600 *A. domesticus*. Of those 600 crickets, half were females and half were males. Half of the female crickets were infested with one larva and the other half was infested with two. We exposed male crickets to the same treatment to test for resource competition in both male and female crickets. Thus, for a single fly’s larvae, 15 females were infested with one larva, 15 females with two, 15 males with one larva, and 15 with two larvae. 150 females were infested with one larva and 150 with two, and 150 males infested with one larva and 150 with two larvae for a total of 600 crickets.

#### Data Analysis

All descriptive and hypotheses testing statistical analyses were conducted in R using R Studio (ver. 2022.02.03 Build 492.pro3). We visually inspected Q-Q plots and used the Shapiro-Wilk Test to assess whether different morphometric data were derived from normally distributed populations. We found that male pronotum width, femur length, mass, and female pronotum width and femur all violated the assumption of normality. Therefore, we used the Spearman’s Rank Correlation to examine the correlation structure between different morphological traits (Fig. 1). Pronotum length was found to be significantly correlated with prontonum width (R^2^ = 0.4419, p <0.0001) and mass (R^2^ = 0.4569, p <0.0001). To avoid the issue of multicollinearity, and following other studies (e.g. Judge and Bonanno, 2008), we chose to use pronotum length as a measure of host cricket size.

Generalized linear mixed-effects models (using glmer in the R package lme4 (ver. 1.1.34)) were applied to investigate the influence of cricket host morphometrics, and level of resource competition on pupation and eclosion success. In these models, cricket size (pronotum length), cricket sex, and level of competition (single larva vs two larvae) were entered as fixed effects, and the fly donor ID was entered as a random effect. Linear regression models were used to determine the relationship between host size (pronotum length) and pupal width (a measure of fly size) and the relationship between pupal width and eclosed F1 mesothorax length.

Statistical comparisons of male and female cricket size, and pupal width in the presence and absence of competition, were accomplished with the Wilcoxon rank sum test using the stats package (ver. 4.1.3) in R.

## Results

### Pupation and Hatching Success Rates

Propagating *Ormia ochracea* in the laboratory can occur with a manual infestation procedure described in Wineriter and Walker (1990) and in Vincent and Bertram (2010b). Larvae were extracted from gravid female *O. ochracea*, and a single larva was placed on top of the membrane underneath the pronotum of a cricket. Within 30 mins, the larva pierced through the membrane and burrowed into the cricket. Larvae were found to emerge from host crickets 6-10 days post parasitization (Fig. 2A), pupated several hours after emergence, and finally eclosed as adult flies 23-27 days later (Fig. 2B). Following previously established manual infestation procedures, the Lee Lab has successfully reared around 50 generations of FL flies over a period of 7 years. Using *Acheta domesticus* as host crickets, in 2023, lab records indicate a success rate (from parasitization to pupation) that varied from 0 to 75%, depending on a combination of factors such as the person performing the manual parasitization procedure, the condition of purchased crickets, and the condition of the larvae extracted from gravid female flies. Based on the combined lab data, our overall pupation success rate was 33% (9,899 pupae collected / 30,171 crickets infested).

**Figure 2.**
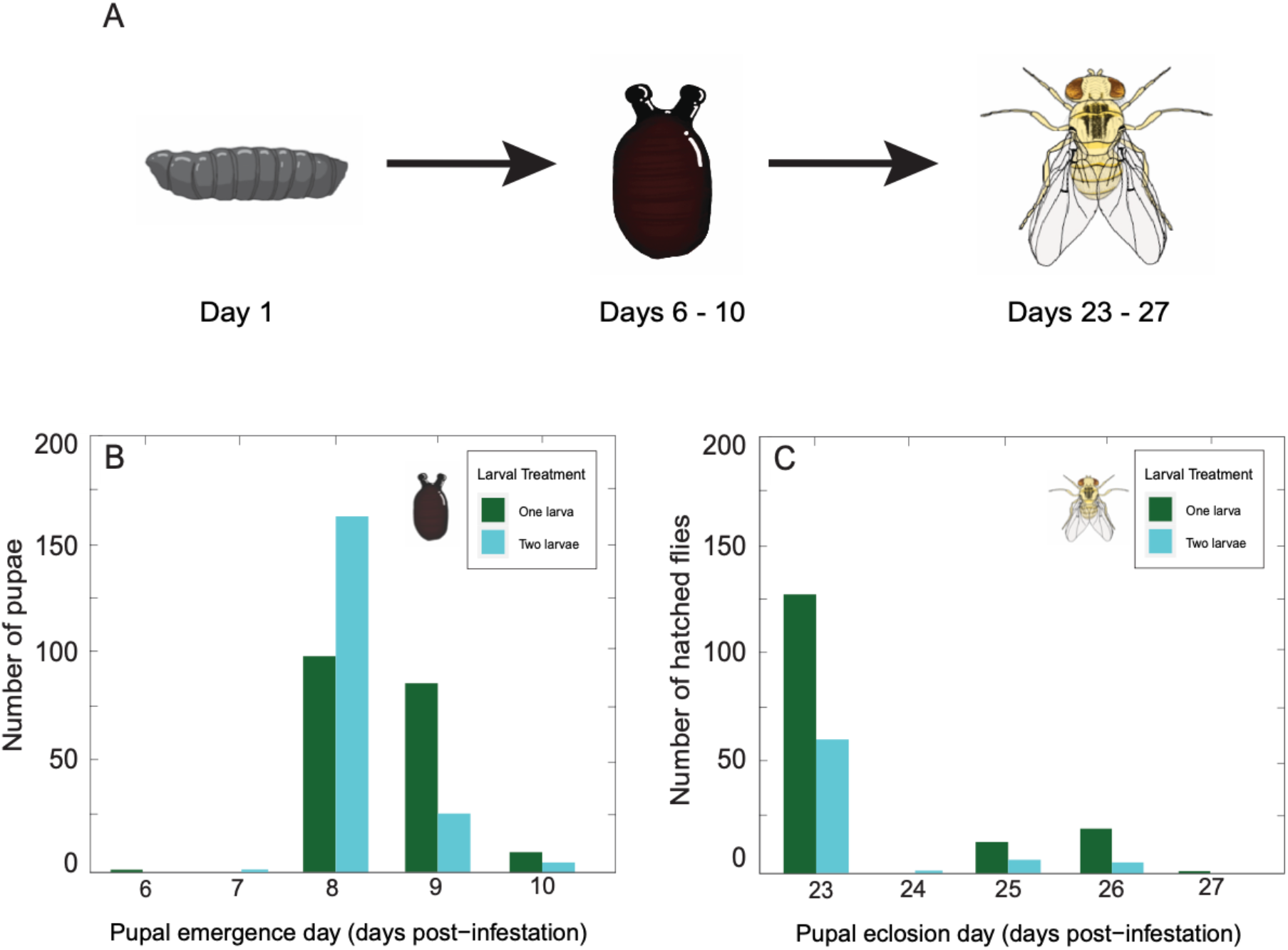
Pupation and eclosion development times. **(A)** First instar plandia, pupae, and, and an adult *Ormia ochracea*. Note: drawings are not to scale. **(B)** Counts of larval emergence and pupation success and **(C)** eclosion success for larvae in developing in the absence (green) or presence of resource competition (cyan) as a function of the number of days post-parasitization.

To examine larval developmental outcomes in detail, we manually infested 600 crickets and maintained them under controlled experimental conditions. We examined pupation success according to two different definitions. First, when pupation success was defined as whether emergence and pupation occurred at all after parasitizing a cricket regardless of the number resulting pupae, neither the level of resource competition in terms of the number of larvae used in parasitization (competition: z = -0.253, p = 0.800), nor cricket sex (cricket sex: z = -1.125, p = 0.260), nor cricket size (pronotum length: z = -0.094, p = 0.925) affected pupation success. Crickets that were parasitized with one larva vs two larvae resulted in similar pupation success rates that were not significantly different from each other (χ^2^ = 0.06, df = 1, p = 0.80) (Fig 3A). When pupation success was defined as the number of larvae that resulted in successful emergence and pupation, we found that neither cricket sex (cricket sex: z = -1.343, p = 0.179), nor cricket size (pronotum length: z = 0.107, p = 0.9144) affected pupation success. Instead, resource competition affected pupation success (competition: z = -3.070, p = 0.002). The proportion of success for larvae that developed in competition (52.67%) was lower than compared to larvae that developed in the absence of competition (63.33%) (Fig 3B).

**Figure 3.**
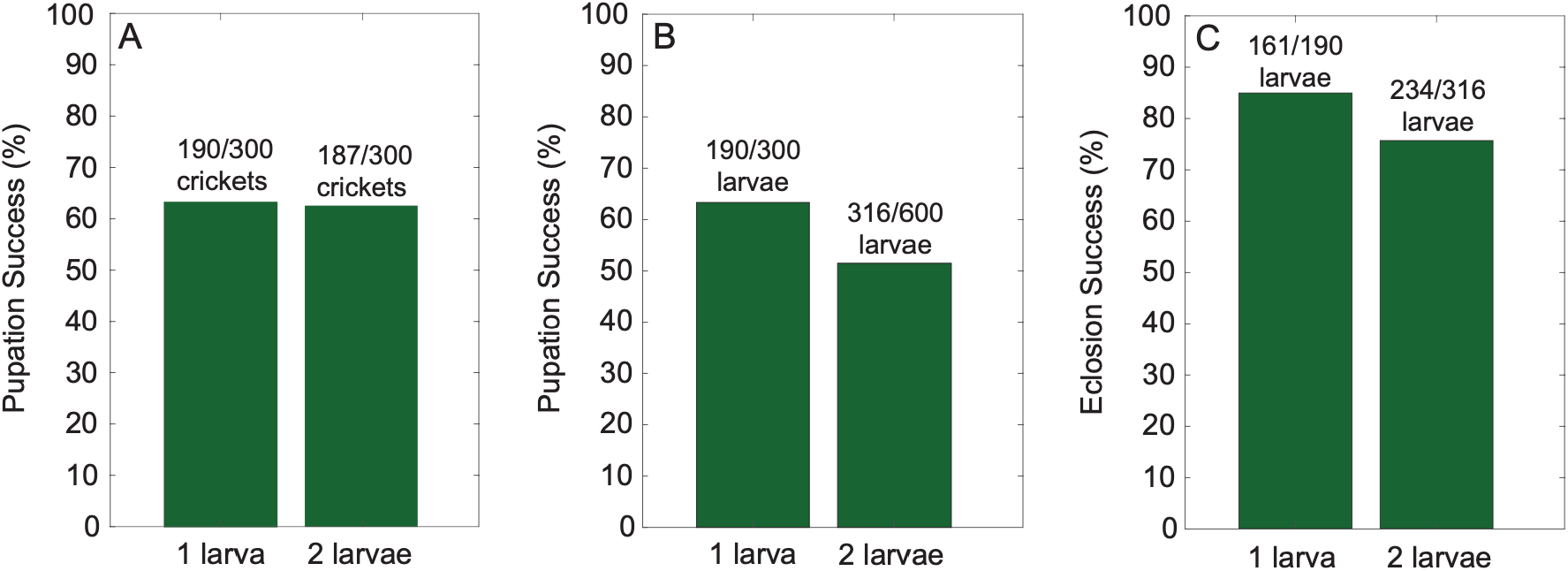
Pupation and eclosion success. **(A)** The percent of larvae that successfully pupated in the absence or presence of resource competition. Pupation success defined as whether emergence and pupation occurred at all after parasitizing a cricket regardless of the number of resulting pupae. **(B)** The percent of larvae that successfully pupated in the absence or presence of resource competition. Pupation success defined as the number of larvae that resulted in successful emergence and pupation. **(C)** The percent of successfully eclosed flies in the absence or presence of resource competition. Counts are expressed above each data bar.

Of the 300 larvae used to parasitize crickets in the absence of resource competition (1 larva per cricket), 63.3% emerged and successfully pupated (190/300) (Fig. 3B, left bar). Whereas among the 600 larvae that experienced resource competition (2 larvae per cricket) only 51.5% successfully pupated (309/600) (Fig. 3B, right bar), resulting in a total success yield of 55.4% across both treatments (499/900). Emergence and pupation started on day 6 post parasitization, with one case in which one larva (0.02% of total successful pupation) successfully emerged and pupated (Fig. 2A). Most larvae emerged and pupated on day 8 post infestation (73.3%, 366/499), while the rest emerged and pupated by day 10 post infestation (day 9: 23.8%, 119/499; day 10: 2.6%, 13/499) (Fig. 2B).

We found that pupal eclosion success depended on the level of resource competition (competition: z = -2.865, p = 0.004, Fig. 3C), but not cricket sex (cricket sex: z = -1.243, p = 0.214), nor cricket size (pronotum length: z = -0.408, p = 0.684). In the absence of competition, 84.74% (161/190) of the pupae successfully eclosed (Fig. 3C, left bar). Eclosion success rate decreased to 74.05% (234/316) when two larvae were competing for resources within the same host cricket (Fig. 3B, right bar). Most of the pupae (83%, 328/395) eclosed 23 days post infestation. In the following days, 0.05% eclosed on day 24 (2/395), 7.34% (29/395) eclosed on day 25, and 9.11% (36/395) eclosed on day 26 after infesting (Fig. 2C).

### The effects of host size on pupal size

The female crickets used in our study were generally larger than male crickets (p < 0.001), with host size ranging from 4.02 to 5.147mm among male crickets, and from 4.159 to 5.257mm among female crickets. In the absence of competition, when crickets were infested with one larva, there was a weak but significant positive relationship between cricket size (pronotum length) and pupal width (adjusted R^2^ = 0.024, F_(1,158)_ = 4.921, p = 0.028, Fig. 4A). This linear regression model revealed that for every millimeter increase in pronotum length, we can expect pupal width to increase by approximately 0.231 millimeters (estimate = 0.231, t = 2.218, p = 0.028).

**Figure 4.**
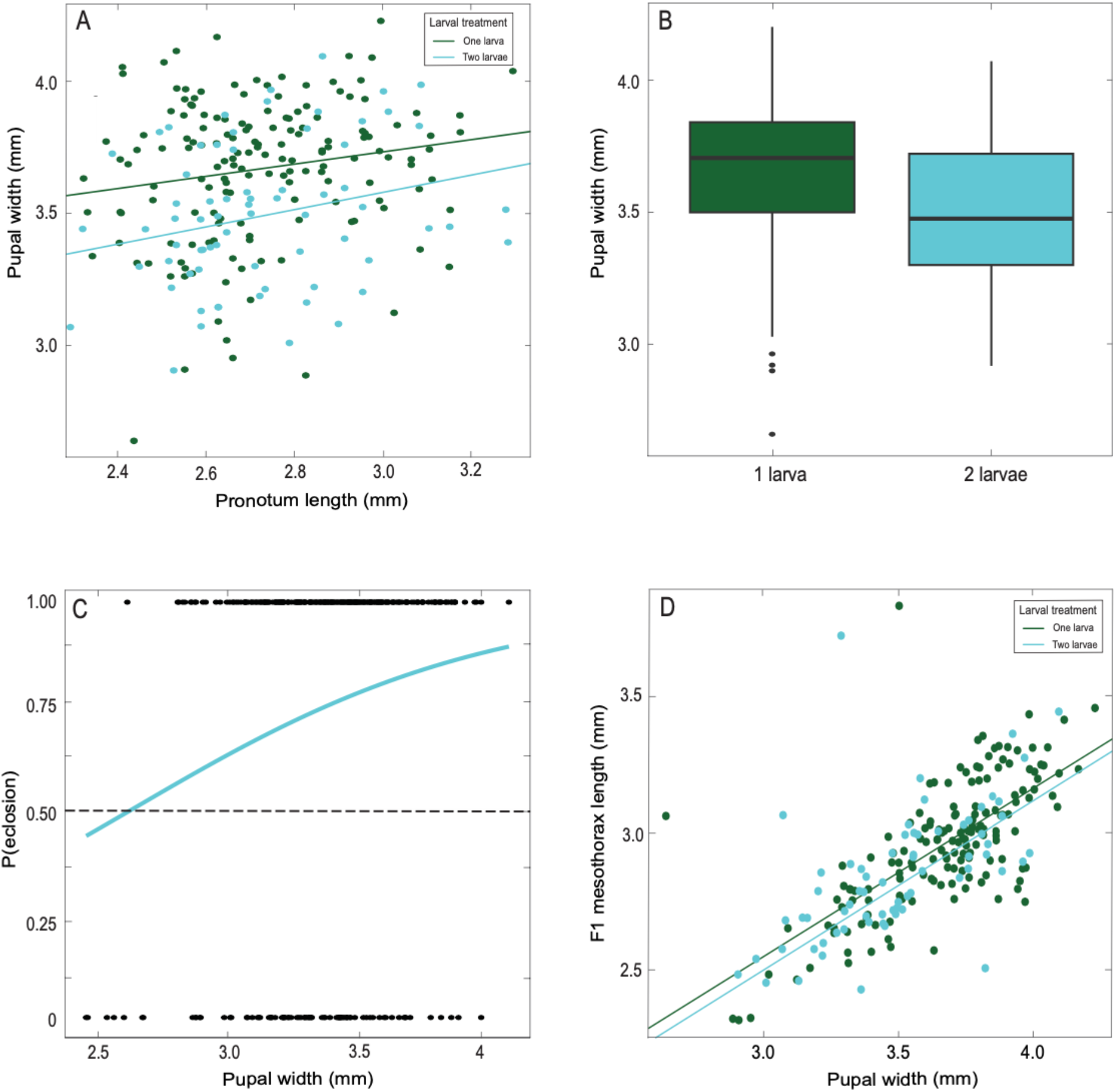
The effects of pupal width on eclosion success. **(A)** Fly pupal width varied with host cricket pronotum length and depended on the level of competition. Best fit lines and data points for 1 larva vs 2 larvae per host cricket indicated in green and cyan, respectively. **(B)** Pupal width as a function of larvae developing in the absence (green) or presence of resource competition (cyan). Box plots depict the first and third quartiles, median values, with the length of the whiskers showing no less (lower) or more (upper) than 1.5 times the interquartile range. **(C)** Probability of eclosion success as a function of pupal width for larvae experiencing resource competition. Fitted line (cyan) from a generalized linear model with a binomial logit link predicts eclosion success. Dotted line depicts chance level (50%) success. Black data points indicate individual successful eclosion events (top), and unsuccessful eclosion events (bottom). **(D)** F1 mesothorax length has a positive relationship with pupal width and did not depend on the level of competition. Best fit lines and data points for 1 larva vs 2 larvae per host indicated in green and cyan, respectively.

### The effects of resource competition on pupal size

Pupal width also depended on the level of competition. Larvae that developed in the absence of competition were significantly larger than those that developed under competition (Wilcoxon rank sum test, W = 7810, p = < 0.0001, Fig. 4B). This relationship was also found to be consistent with the eclosed adult flies (Wilcoxon rank sum test, W = 7591.5, p = 0.0000456, Fig. S1). When crickets were infested with two larvae, we also found a weak but significant relationship between cricket host size and pupal width (adjusted R^2^ = 0.048, F_(1, 69)_ = 4.531, p = 0.037, Fig. 4A). In this case, for every unit increase in pronotum length, we can expect pupal width to increase by 0.327 units (estimate = 0.3269, t = 2.129, p = 0.037). Comparing the two best fit lines, pupal width was greater in the absence of competition across all pronotum lengths (Fig. 4A). We also found that pupal width predicted eclosion success (p = 0.00147, Fig. 4C), and also varied significantly with eclosed adult fly size (mesothorax length) (1 larva: adjusted R^2^ = 0.487, F_(1, 158)_ = 149.9, p = < 0.0001, 2 larvae: adjusted R^2^ = 0.261, F_(1, 69)_ = 25.7, p = < 0.0001, Fig. 4D). At a pupal width of 2.62 mm, 50% of the pupae can be expected to eclose successfully (Fig. 4C).

## Discussion

In manual parasitizations with commercially acquired *Acheta domesticus* as host crickets, we found that neither host size, nor cricket sex, affected pupation success or eclosion success. However, resource competition negatively impacted both pupation and eclosion success (Fig. 3B,C) and the size of developing pupae (Fig. 4A,B). Larvae that developed in the absence of resource competition generally resulted in larger pupae than compared to pupae that developed under resource competition (Fig. 4B). Therefore, we found some experimental evidence in support of our hypothesis that resource competition can negatively impact larval developmental outcomes in terms of pupal size, and pupation and eclosion success.

*Ormia ochracea* larvae can develop in a range of natural host cricket species that vary depending on geographic location (Gray et al., 2019, 2007; Sakaguchi and Gray, 2011; Walker, 1993; Walker and Wineriter, 1991). In the laboratory, manual parasitization experiments have established that *O. ochracea* larvae can successfully develop in other cricket species that are not known to be their natural hosts, but pupation success can be lower than compared to those developing in natural hosts (Adamo et al., 1995a; Broder et al., 2023; Thomson et al., 2012; Wineriter and Walker, 1990). For example, 54 - 61% of manually parasitized *Gryllus texensis* (natural host) resulted in successful larval emergence and pupation, while only 32 - 35% (55 used in this study) of manually parasitized *Gryllus assimilis* resulted in successful larval emergence and pupation (Thomson et al., 2012). In Vincent and Bertram (2009), manual parasitization of adult *G. texensis* resulted in an average of 78% pupation rate, and they found similar pupation success rates in manually parasitizing juvenile *G. texensis*. Interestingly, Adamo et al. (1995b) had a much higher emergence and pupation success rate of 90% (27/30 crickets in study) while manually parasitizing *G. texensis* (previously identified as *Gryllus integer* in the paper). Wineriter and Walker (1990) report in their abstract that “commercially reared *Acheta domesticus* tested as hosts were less satisfactory”, but no data is provided in the paper to clarify the nature of this statement. The overall pupation success rate of 55.4% in this study supports the idea that the unnatural cricket host *A. domesticus* may provide similar pupae yields relative to natural host cricket species (e.g. compared to the Thomson et. al. 2012 study). Eclosion success in *A. domesticus* depended on the level of resource competition. When crickets were singly parasitized, 84.74% successfully eclosed. This was significantly better than singly parasitized natural hosts that eclosed at rates of 32 - 45% success (Thomson et al., 2012). Eclosion success rate dropped to 74.05% when crickets were parasitized with two larvae. Adamo et al. (1995b) did not report on eclosion success rates in terms of percentage success, but they found that the number of flies eclosed peaked when natural host crickets (*G. rubens* and *G. texensis*) were parasitized with four larvae. The number of flies eclosed was reduced with further increases in the number of larvae used for parasitizing a single host cricket (Adamo et al., 1995b). Despite male crickets generally being smaller than female crickets, in agreement with Thomson et al. (2012), we did not find cricket sex to significantly affect pupation or eclosion success.

By contrast, resource competition affected the size of the pupae, and eclosion success, which is also expected to affect adult fly size. Larvae that experienced resource competition were smaller (Fig. 4B) and less likely to successfully eclose (Fig. 4C). These results are consistent with findings presented in Adamo et al. (1995b), which demonstrated that pupal size, number of pupae, and number of hatched adults, all decreased with increasing larval numbers per cricket in both *G. rubens* and *G. texensis*. In *O. ochracea*, the number of planidia that gravid females carry can range from less than 65 to more than 500 (Wineriter and Walker, 1990). While not directly examined in *O. ochracea*, fecundity in other tachinids vary with fly size (Ho et al., 2011; Lauziere et al., 2001; Nakamura, 1995; Reitz and Adler, 1995). Presumably, fly size in *O. ochracea* will also affect fecundity and overall reproductive fitness.

In the field, superparasitism (multiple larvae of distinct size class within the same host cricket) occurs at a low rate (Adamo et al., 1995b). As muscle tissue is spared during the first few days of larval development, male crickets maintain the ability to produce calling songs and thus can still attract eavesdropping flies for several days after being parasitized (Adamo et al., 1995a). According to this situation, early developing larva will emerge and result in cricket death before subsequent larvae are able to successfully emerge (Adamo et al., 1995b). Thus, it pays for *O. ochracea* to avoid superparasitism, but behavioral experiments demonstrate that *O. ochracea* are unable to distinguish between unparasitized versus parasitized crickets (host discrimination) (Adamo et al., 1995b). The most optimal reproductive strategy for *O. ochracea* is to deposit a sufficient number of larvae to guarantee at least one larva is established in a suitable host cricket. This will mitigate the effects of resource competition in affecting the size of the pupae, which seems to also determine eclosion success.

Compared to *Gryllus* and *Teleogryllus* field cricket species, *A. domesticus* tends to be smaller in size. This size difference, or potential differences in cricket immune function (e.g. Sikkink et al., (2020)), may contribute to differences in pupation and/or eclosion success. Across the continental US, *O. ochracea* are known to parasitize at least 17 host species (Gray et al., 2019), and are found to be behaviorally specialized in parasitizing one or a few species within a geographic location (Gray et al., 2007). In Hawaii, where the preferred host cricket *Teleogryllus oceanicus* continue to evolve in response to selection imposed by *O. ochracea*, some crickets have lost the ability to produce calling songs (Rayner et al., 2019; Zuk et al., 2006), while others have been rapidly evolving more cryptic calling songs with frequency content that differs from ancestral *Teleogryllus oceanicus* calling songs (Broder et al., 2022; Gallagher et al., 2022; Tinghitella et al., 2021, 2018). These changes to cricket calling songs may have led to the switching to alternative host species (Broder et al., 2023). At present, it is unknown what allows a field cricket species to be suitable for the development of *O. ochracea* larvae.

As larval development generally depends on the availability of host resources (Godfray, 1994; Harvey et al., 2013), we expect our findings on resource competition and larval developmental outcomes to be generalizable across other different species of *Ormia*. It would be of interest to determine if use of *Acheta domesticus* can be viable and yield similar development outcomes for other acoustic parasitoid flies (Lakes-Harlan and Lehmann, 2015; Lehmann, 2003) that parasitize cicadas (e.g. Schniederkotter and Lakes-Harlan, 2004) or katydids (Rogers and Beckers, 2023).

## Conclusions

Our results demonstrate that resource competition can negatively affect the developmental outcome of *O. ochracea*. These results also show the feasibility of using *Acheta domesticus* in propagating colonies of *O. ochracea* in the laboratory. Ideally, recently molted adult *Acheta domesticus* should be used as host crickets. To allow for the largest possible pupal width and fly size, and to maximize pupation and eclosion success, each host cricket should only be parasitized with a single larva. Stable colonies of *O. ochracea* can be sustainable across multiple generations; we manually parasitize approximately 4000 crickets per month, each with a single larva to maximize fly size and eclosion success. Parasitized crickets are housed in a communal setting (100 crickets in a 16-quart Sterilite storage container). This ensures approximately 1300 pupae that can be divided evenly among three 20 quart Sterilite™ fly population containers for eclosion. This fly density ensures that matings will occur to provide a good supply of gravid female flies for colony propagation and experimentation.

## Acknowledgements

We would like to thank The Bug Company (Ham Lake, MN), especially their staff for providing us with a constant reliable source of *Acheta domesticus* that supports research activities in the Lee Lab at St. Olaf College. We thank past and present members of Lee Lab for help with animal care and propagating fly colonies. Funding was provided by a National Science Foundation CAREER grant to NL (IOS 2144831) and support from the St. Olaf Collaborative Undergraduate Research and Inquiry (CURI) Program and the St. Olaf TRIO McNair Program. BL was funded by a UK-Canada Globalink Doctoral Exchange Scheme grant from the UK Research and Innovation (UKRI) council in partnership with Mitacs (Canadian not-for-profit organization). ACM was supported by NSERC Canada grant #2020-05946.

